# Reduction of Pollen Number and Anther Length in Bread Wheat Studied by a Nested Association Mapping Population

**DOI:** 10.64898/2026.05.22.727104

**Authors:** Naoto-Benjamin Hamaya, Hiroyuki Kakui, Moeko Okada, Jilu Nie, Katharina Jung, Miyuki Nitta, Thomas Wicker, Beat Keller, Shuhei Nasuda, Kentaro K. Shimizu

## Abstract

The number of pollen grains, which carry male gametes in seed plants, has attracted interest in genetics, evolution, and breeding. Rapid evolutionary reductions in pollen number and anther length were reported in selfing species as well as domesticated species, although this poses a challenge for hybrid breeding. Here, we studied the variation of pollen number and anther length of the hexaploid bread wheat (*Triticum aestivum*) by employing a quick pollen counting method. Pollen numbers in cultivars were lower than those in landraces among 54 lines of diverse geographic origins. Using the year of registration of traditional and modern cultivars, we found a reduction in pollen number over the past 150 years. We detected high heritability and variation among Asian landraces and cultivars. Thus, we conducted QTL mapping of pollen number as well as of anther length using nested association mapping lines in which Norin 61 was the common parent. Genomic loci encompassing Green Revolution genes (*Rht-B1*, *Rht-D1,* and *Ppd-D1*) showed significant effects on pollen number and anther length, but their contributions were relatively minor. Although anther length has often been used as a proxy for pollen number in bread wheat, our data showed that their correlations are not necessarily high. Interestingly, we identified a new QTL of pollen number that was not detected by measuring anther length, and, vice versa, a new QTL specific to anther length. These data suggest that pollen number has reduced rapidly in bread wheat but can be modified using the genetic diversity of landraces.

**Significance statement:** We found that modern cultivars of bread wheat have reduced pollen number and shorter anther length, which are common in domesticated species but can be a challenge for hybrid breeding. Using underutilized Asian landraces and cultivars, we reported that new quantitative trait loci as well as loci used in the Green Revolution, are responsible for the traits, which can be employed to increase pollen numbers.

## Introduction

Anther and pollen development are crucial for successful plant reproduction. Following the release of pollen grains from the anther and their deposition at the top of the pistil, the pollen tube grows to the ovule to deliver two sperm cells, initiating double fertilization (Higashiyama and Takeuchi, 2015; Shimizu and Okada, 2000). Numerous studies have identified genes regulating anther cell differentiation and pollen formation ranging from model species i.e., *Arabidopsis thaliana*, to a lesser extent to crop species i.e., maize and rice (Gómez *et al*., 2015; Marchant and Walbot, 2022). Nevertheless, natural variation in pollen number and anther length is still poorly understood (Guo *et al*., 2023; Tsuchimatsu *et al*., 2020). Variation of male gamete number, i.e., pollen number in plants and sperm number in animals, has attracted a broad interest in evolutionary biology, agriculture, and medicine (Darwin, 1876; Harvey and May, 1989; Lüpold and Fitzpatrick, 2015; Shimizu and Tsuchimatsu, 2015; Sicard and Lenhard, 2011). Evolutionary analysis on sex allocation theory suggested that reduced pollen number can be advantageous to allocate less resource to male reproductive success when the outcrossing rate is low (Charnov, 1982; Shimizu and Tsuchimatsu, 2015). The reduced number of pollen grains is a hallmark of the selfing syndrome, or a reduction in floral traits such as reduced anther extrusion in predominantly selfing species (Shimizu and Tsuchimatsu, 2015; Tsuchimatsu and Fujii, 2022). The reduced pollen number in selfing species compared to closely related outcrossing species is often represented by a reduced pollen/ovule ratio (Cruden, 2000). The ratio is identical to the pollen number in a single floret (flower) in grasses because a grass floret has a single ovule. Selfing syndrome is commonly found both in natural and crop species. Domesticated rice was reported to show a smaller anther length, which is typically used as a proxy of pollen number, in accordance with low outcrossing rate (Oka and Morishima, 1967).

To study the genetic basis of pollen number as a quantitative traits, genome-wide association study (GWAS) and QTL mapping using bi-parental populations have been employed. Recent studies started to reveal the molecular basis of the variation of pollen numbers (Kakui *et al*., 2022; Tsuchimatsu *et al*., 2020). A GWAS in *A. thaliana* revealed the gene *REDUCED POLLEN NUMBER 1* (*RDP1*), which explained approximately 20% of the variation in pollen number per flower between accessions in the study. *RDP1* was the first gene identified to affect the natural variation of the number of male gametes and notably showed signatures of selection for a reduced pollen number (Tsuchimatsu *et al*., 2020). *RDP1* is a putative ribosome biogenesis gene (Tsuchimatsu *et al*., 2020), which is supported by the observed upregulation of ribosome genes and downregulation of pollen development genes in the *rdp1-3* mutant (Kakui *et al*., 2022). Still, the underlying genetic mechanism remains elusive. Recently, two genes have been identified as negative regulators of pollen number in maize: *ZmCCT10* and *ZmRPN1* (Guo *et al*., 2023; B., Li *et al*., 2022). *ZmCCT10* was first characterized as photoperiod-sensitive negative regulator of flowering which encodes a *CCT* (*CO, CO-LIKE* and *TIMING OF CAB1*) domain transcription factor (Liu *et al*., 2020; Yang *et al*., 2013). Loss of *ZmCCT10* function increases pollen number (Li *et al*., 2022). *ZmRPN1*, which was identified through a GWAS, functions as another negative regulator of pollen number by supporting the localization of *ZmMSP1*, a regulator of cell differentiation in somatic tapetal and reproductive lineages (Guo *et al*., 2023; Nonomura *et al*., 2003; Zhao *et al*., 2002).

Hybrid breeding can achieve higher yields in many crop species by masking the effects of deleterious variants in F_1_ hybrids (ter Steeg *et al*., 2022; Whitford *et al*., 2013), and is also effective in polyploid species such as bread wheat (Revell *et al*., 2025; Whitford *et al*., 2013). To obtain F_1_ seeds, a paternal pollen donor cultivar should shed an adequate amount of pollen grains in order to pollinate a neighboring male-sterile cultivar. However, domesticated species such as rice were reported to show smaller anther lengths (Oka and Morishima 1967). Low fertility is often reported in modern agricultural cultivars, potentially due to accumulation of deleterious mutations associated with genetic bottlenecks during domestication and subsequent breeding (Jiang *et al*., 2024). In bread wheat, despite the presence of duplicated homeologs, purifying selection on deleterious variants are detected, which provide the molecular basis of heterosis (Halstead-Nussloch *et al*., 2026). Facilitating hybrid breeding is a major goal in wheat breeding, but traits associated with the selfing syndrome have been a barrier for hybrid breeding in predominantly selfing wheat (Burgarella *et al*., 2024; Okada *et al*., 2019; Whitford *et al*., 2013). To achieve high pollen emission of paternal plants, a high number of pollen grains (or large anthers as a proxy) is necessary to be combined with other beneficial floral traits to promote outcrossing such as anther extrusions and increased height (Boeven *et al*., 2016; De Vries, 1974; Langer *et al*., 2014; Okada *et al*., 2019; Rohde *et al*., 2025). Several studies showed that genes introduced during the Green Revolution in the 1960s (Calderini and Slafer 1998) also had effects on traits related to the efficiency of hybrid breeding. Pleiotropic effects such as the reduction of anther length were shown for the genomic regions encompassing *Rht-B1b* (*Rht1*), *Rht-D1b* (*Rht2*), *Rht-B1c* (*Rht3*), and photoperiod insensitive *Ppd-B1a* and *Ppd-D1a* (Okada *et al*., 2019; Schierenbeck *et al*., 2024). It is possible that these *Rht* and *Ppd* alleles affect pollen number assuming the positive correlation between pollen number and anther length (Schierenbeck *et al*., 2024).

To study the genetic basis of quantitative traits, nested association mapping (NAM) populations have been initially developed in maize to integrate the advantages of genome-wide association studies and bi-parental QTL mapping (Buckler *et al*., 2009; Gage *et al*., 2020; Yu *et al*., 2008). In NAM populations, a single common parental line is crossed to multiple lines to cover a broader genetic variation within a species. Then, recombinant inbred lines are generated from each cross to induce recombination events. Subsequently, lines are genotyped for the construction of linkage maps and for the detection of QTLs. In wheat, the usage of NAM lines was facilitated by the chromosome-scale genome assemblies of diverse cultivars including the experimental cultivar Chinese Spring (CS) (IWGSC *et al*., 2018) and ten globally representative cultivars of the 10+ wheat genome project (Shimizu *et al*., 2021; Walkowiak *et al*., 2020; White *et al*., 2025). Recently, this progress in genome assemblies was utilized to study disease resistance in an wheat NAM population consisting of Asian landraces and modern cultivars (Jung *et al*., 2025), which are underrepresented in modern breeding (Balfourier *et al*., 2019).

An obstacle in studying loci affecting the natural variation of pollen number is that phenotyping of pollen number represents a large workload when done classically by microscopy observations. Previous studies used anther length as a proxy for pollen number to overcome this issue, since relatively high correlations between the two traits are reported (De Vries, 1974; Milohnic *et al*., 1976; Nguyen *et al*., 2015). To overcome the hurdle of slow microscopy observations, we developed a method that can efficiently and directly measure the pollen number of wheat and *Arabidopsis* in a large mapping population by using a cell counter (Kakui *et al*., 2020a; Kakui *et al*., 2020b; Tsuchimatsu *et al*., 2020). In this study, we examined pollen number and anther length to elucidate how breeding shaped the two traits from landraces to modern cultivars. We first measured pollen number and anther length of 54 lines of bread wheat consisting of landraces and cultivars from a diverse geographic origin. We examined the temporal change in these traits in the past 200 years and further investigated pollen number of 25 lines in controlled chamber environments under control and moderate heat stress conditions during reproductive development. Based on the high heritability of pollen number and anther length, we conducted multiparent QTL mapping on these traits in a NAM population consisting of 1057 NAM lines. Finally, we genotyped the 54 lines to examine the contribution of genomic regions encompassing green revolution genes.

## Material and Methods

### Plant materials

#### 1) Landraces and cultivars

We assembled a small subset of a wheat diversity panel composed of 54 wheat lines including landraces and cultivars to study differences in pollen-related traits (Suppl. Table 1). Landraces are defined as wheat lines that spread globally after domestication and adapted to the different local environments along human migration routes, while cultivars are considered to originate from breeding programs that used landraces as a source to systematically improve the crop (Balfourier et al. 2019). The lines included 25 parental lines of the National BioResource Project-Wheat (NBRP-Wheat) nested association mapping (NAM) population composed of Asian landraces and cultivars, 13 cultivars from the 10+ wheat genome project (Walkowiak et al. 2020) (the spelt variety PI190962 was not included and Norin 61 was already part of the 25 parental lines of the NAM population), as well as 9 landraces and 7 cultivars used in Balfourier et al. (2019) to cover a broad range of genetic diversity. The 31 cultivars used in this study were further divided into the sub-categories, of 12 traditional and 19 modern cultivars. Traditional cultivars are wheat lines that were registered before 1960, while modern cultivars were released after 1960 (Balfourier *et al*., 2019). We obtained wheat lines through the John Innes Centre (https://www.seedstor.ac.uk/index.php), the INRA Small Grain Cereals Biological Resources Centre (INRA) (https://urgi.versailles.inra.fr/siregal/siregal/grc.do) and the National BioResource Project-Komugi (NBRP-Wheat, https://shigen.nig.ac.jp/wheat/komugi/). Arina*LrFor* was donated by Prof. Beat Keller, University of Zurich (Suppl. Table 1).

#### 2) NAM population

The NBRP-Wheat NAM population consists of 25 Asian parental lines including landraces and cultivars (Table 1). The parental lines were selected from the core collection of hexaploid wheat from the Japanese gene bank NBRP-Wheat to cover a wide genetic diversity in Asia (Takenaka *et al*., 2018). The lines originate from Japan, China, Nepal and Pakistan. The line NP4 was previously classified to be of Bhutanese origin in Takenaka et al. 2018 and has been revised to be of Nepalese origin. The common male parent Norin 61 (Tsujimoto, 2021) was crossed with each of the 24 female parental lines, and selfed until F_7_ generation to form the Recombinant Inbred Lines (RIL) (Suppl. Figure 1). We selected 13 RIL families containing 50-160 individuals based on parental phenotypes, such as anther length and pollen number, but also disease resistance (Table 1). All of the seeds of 25 NAM parental lines and 13 RIL families were obtained from NBRP-Wheat.

**Table 1:**
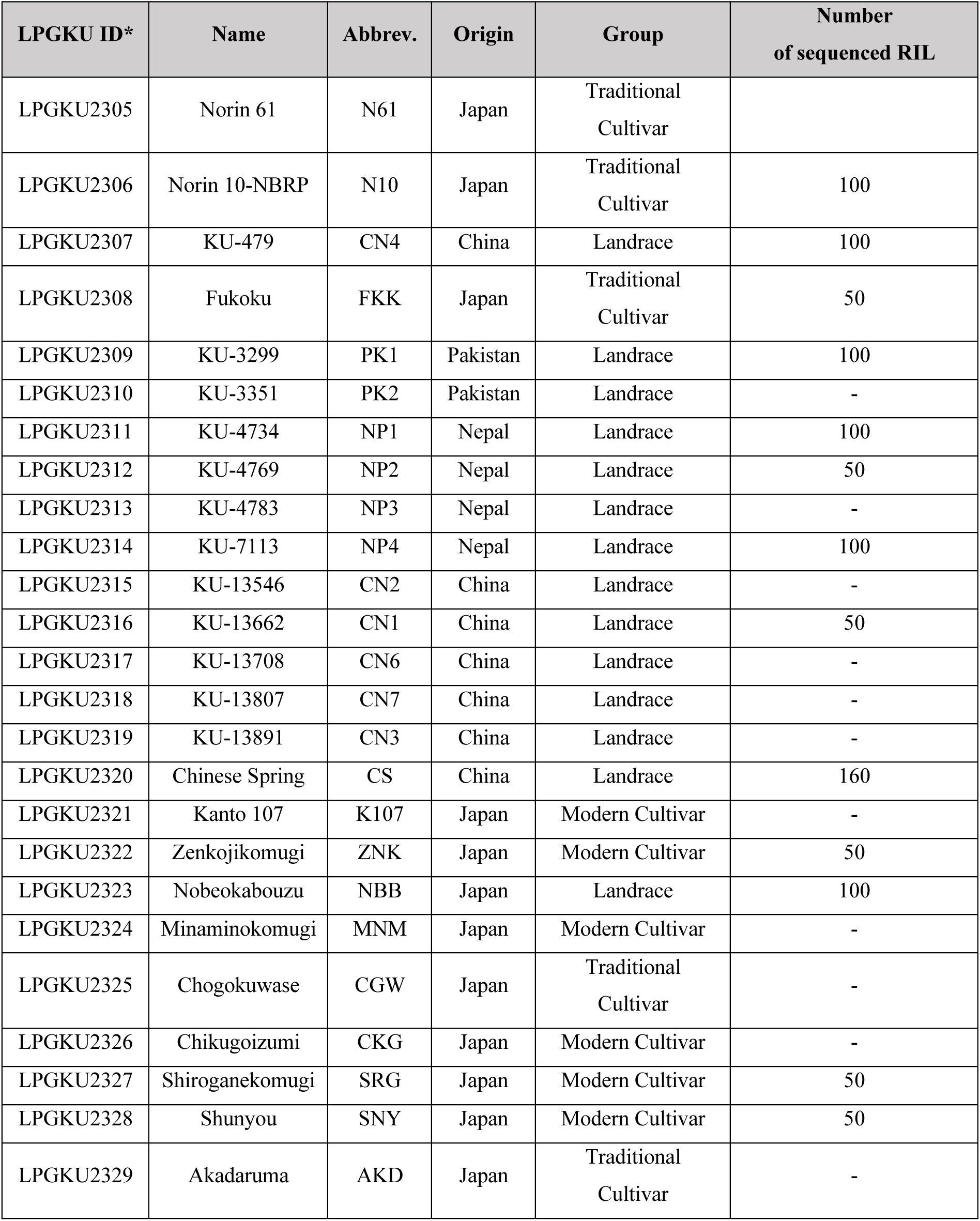
Information about the 25 NAM parental lines. Norin 61 is the male parental line. *LPGKU (Laboratory of Plant Genetics, Kyoto University) ID is accessible through NBRP-Wheat.

### Plant growth conditions

#### 1) Growing subset of wheat diversity panel in foil greenhouse

Twenty seeds of the selected 54 wheat lines were sown directly into soil in October 2021 in a foil greenhouse on Campus Irchel at the University of Zurich, Switzerland. To ensure a uniform germination in the second year, seeds were sterilized by rinsing them with 70% Ethanol and then left in a 0.5% Javel water solution for 30 min. Twelve germinated seeds were then transplanted to a strip of twelve organic planting pots 4.5 x 4.5 cm (Jiffystrips, Jiffy) filled with soil (Einheitserde Classic, Einheitserde) and grown for two weeks at short day conditions (18°C / 16°C; 8 h light / 16 h darkness). Followingly, the strip was separated into single units and eight seedlings per genotype were directly transplanted into soil in the foil greenhouse. After germination, eight seedlings were kept, and excess plants were removed. Plants were treated with Amistar Xtra (Syngenta Agro AG, Switzerland) to prevent the spread of *Blumeria graminis* f.sp. *tritici*, since they were grown in a semi-natural environment. During the growth period in Spring, the plants received solid fertilizer (Polydor, Maag) and approximately every three weeks liquid fertilizer (Wuxal Universaldünger, Hauert, Switzerland).

#### 2) NAM parental lines in growth chamber

We grew the 25 NAM parental lines two times in growth chambers. The goal of the first experiment was to assess the variation in pollen number. Hence, we grew the plants under control conditions in a growth chamber (22°C / 20°C; 16 h light / 8 h darkness). Seeds were sterilized the same way as in the previous Section 1). First, seeds were then placed in water overnight and transferred to petri dishes containing two layers of round filter paper (90mm diameter, Macherey-Nagel, Germany) soaked with water. Petri dishes were then closed with Bemis Parafilm M (Fisher Scientific, Germany), wrapped with aluminum foil and left for four days at 4°C. Afterwards, aluminum foil was removed, and the petri dishes were brought for germination to a growth chamber (18°C / 16°C; 8 h light / 16 h darkness). After germination, twelve seedlings of each line were transplanted to strips of twelve organic pots 4.5 x 4.5 cm (Jiffystrips, Jiffy, Netherlands) filled with soil (Einheitserde Classic, Einheitserde, Germany) and grown for two weeks in the growth chamber. The seedlings were then vernalized for 3 months at 4°C under short-day conditions. After vernalization, four seedlings per line were transplanted into 11 x 11 x 20 (2 liters) pots filled with soil (Einheitserde Classic, Einheitserde) and solid fertilizer (Polydor, Maag, Switzerland) added to the top layer of the soil. Liquid fertilizer was added every 2-3 weeks (Wuxal Universaldünger, Hauert, Switzerland). Japanese cultivar CGW was excluded from the analysis, as sample number was low (*n* = 2) due to missed sampling caused by very early flowering phenotype.

In the second experiment, we assessed the effect of moderate heat stress from the mid boot stage until flowering on pollen number and pollen diameter by comparing plants grown under control conditions and plants subjected to heat stress. First, we sterilized and germinated NAM parental line seeds with eight replicates according to the protocol described above. In the beginning, all plants were grown under long day conditions (22°C / 20°C; 16 h light / 8 h darkness). Four out of eight replicates of each genotype were separately moved to a chamber with moderate heat conditions (32°C / 26°C; 16 h light / 8 h darkness). We moved the plants when they reached the mid boot stage (BBCH 43) in the second tiller (Lancashire *et al*., 1991). We focused on the second tiller, since we considered it to be less likely to miss the time point of booting compared to the first tiller. The plants stayed in the heat stress conditions from mid boot stage (BBCH 43) until anthers turned yellow (BBCH 59-61) according to the BBCH-scale (cereals) (Lancashire *et al*., 1991). We assessed pollen number and diameter for all four replicates of each genotype that stayed in control conditions and the four replicates subjected to heat stress.

#### 3) Growth of the NAM population in the field

Based on parental phenotypes, 13 families with a total of 1060 RIL were selected (Suppl. Table 2). The RIL seeds were sown in December 2020 in cell trays (Meiwa (Cat. No. 53930), Japan) and grown in a greenhouse until they were transplanted to the field in January 2021 at the experimental farm of Kyoto University in Kyoto, Japan. Ten individuals were grown per genotype. Out of 1060 RIL, 1057 were analyzed as described in the section anther sampling.

### Anther sampling

Anthers were sampled from the middle of the spike by opening the florets with forceps at growth stages 59-61 according to the BBCH-scale (cereals) (Lancashire *et al*., 1991). Anthers were mature at the point of sampling but still closed. We sampled three anthers per floret for the lines grown in the foil house, while one anther per floret was used for the analysis of NAM parental lines grown in the two growth chamber experiments. Samples were dried overnight at 60°C and then stored at -20 °C until they were used for pollen counting.

Sample numbers per line varied between experiments. In the foil house experiments, four samples per line were collected for the first season and eight in the second. In the first growth chamber experiment, all lines had over 14 samples, except for CGW (n=2), which was therefore excluded from subsequent analyses. For the second growth chamber experiment 20 samples per line were collected under both control and heat stress conditions.

For the selected RILs, two spikes were cut for each RIL in the field from plants at growth stages 59-61 according to the BBCH-scale (cereals) (Lancashire *et al*., 1991). The spikes were placed into a 15ml tube stand placed in a box filled with water and brought to the laboratory. For each of the spikes, six mature and closed anthers were then sampled from two florets located in the middle of the spike and placed into a 1.5ml tube. The samples of three genotypes of the 1060 RIL from the mapping population families (KU-N11-49, KU-N15-67, KU-N22-33) were lost during sampling, resulting in 1057 lines used for subsequent analyses.

### Measuring anther length

Pictures of anthers, that were sampled in Zurich in the foil greenhouse during 2023, were taken using a stereo microscope (SZX12, Olympus, Japan) with a monochrome camera (XM10, Olympus, Japan). Anther photos of 1057 RIL were taken by a stereo microscope (SZX7, Olympus) with CCD camera (DP22, Olympus). Anthers were then measured with the image processing software Fiji (Schindelin *et al*., 2012).

### Pollen counting

The number of pollen grains were analyzed based on protocols of Kakui *et al*. (2020a) and Kakui *et al*. (2020b). The pollen grains were released by adding 100 µl ddH_2_O into the 1.5 ml tubes in which the anthers were stored and crushing the anthers with a pestle. The resulting pollen suspension was filtered by spinning it down on a self-made 20 µm filter column for 10 sec at 2000 x *g* (Kakui *et al*., 2020b). The pollen grains were then transferred into the CASYcup (OLS OMNI Life Science, Germany) measurement tube by resuspending 200 µl CASYton (OLS OMNI Life Science) taken from the CASYcup and pipetted on the filter column. This resuspension step was repeated two times. Followingly, the number of pollen grains and pollen diameter were measured by using a cell counter (CASY Model TT, OLS OMNI Life Science) (Kakui *et al*., 2020a). Pollen numbers are measured per anther for all experiments conducted in the growth chambers, while pollen number per floret was measured for experiments performed in the foil house and in the field.

### Group comparison and correlation between landraces and cultivars

We compared pollen numbers, anther length, and pollen diameter between landraces and cultivars by running non-parametric Kruskal-Wallis rank sum test. When Kruskal–Wallis test indicated significant differences, pairwise comparisons were performed using Dunn’s post-hoc test with Bonferroni correction for multiple testing. Dunn’s post-hoc test was performed by using the R package ‘FSA’ (Ogle *et al*., 2025). Furthermore, we calculated Pearson’s correlation coefficient *r* to evaluate significant associations among traits.

### Calculation of broad-sense heritability

We calculated broad-sense heritability for anther length, pollen number and pollen diameter using the NAM parental lines grown in the growth chamber, all lines grown in the foil greenhouse in season 2022 and 2023. First, we built a linear mixed-effect model for each trait in each environment with genotype and replicate as random effects using the R package *lme4* (Bates *et al*., 2015). Variance components were then extracted and broad-sense heritability (*H^2^*) was calculated based on the formula of (Hallauer, 2010)). Replicate number *r* was calculated by the harmonic mean of all replicates per genotype.

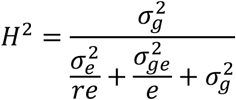

### Calculation of best linear unbiased prediction (BLUP) values

We calculated BLUP values for anther length, pollen number and pollen diameter for the NAM RIL, by building linear mixed effect models using the R package *lme4* (Bates *et al*., 2015). We selected the best fitting model based on Akaike information criterion (AIC) values. The residuals of the best model were tested for a normal distribution using Kolmogorov-Smirnov test. Broad-sense heritability for the NAM RILs was calculated as described above. BLUP values for pollen diameter were not calculated because model assumptions of linear-mixed effect models were violated. The calculated BLUP values were used as input for multiparent QTL mapping.

### Sequencing data and QTL mapping

All 1060 RIL were sequenced with the whole-genome amplicon sequencing technique of GRAS-Di (Enoki and Takeuchi, 2018; Hosoya *et al*., 2019). GRAS-Di raw data is available at DDBJ under accession number DRA019457. SNPs were called on Norin 61 as reference sequence (GCA_904066035.1), since it is the common male parent in the NAM population and due to its high-quality reference genome (Shimizu *et al*., 2021; Walkowiak *et al*., 2020). 6467 markers were used to create the consensus map for QTL mapping. Refer to Nie (2025) for the full workflow and Jung et al. (2025) for a short summary. QTL mapping was performed using the R package *statgenMPP* version 1.0.4 (W., Li *et al*., 2022). First, identity by descent (IBD) probabilities were calculated using the *calcIBDMPP* function with *F7* specified as population type. The evaluation distance was 1 cM. Second, a significance threshold for the QTL mapping was calculated using a permutation test on single-marker QTL mapping without any kinship matrix due to increased computation time and the small effect that it has on the final mapping (Jung *et al*., 2025). The permutation test was run by randomly shuffling the phenotype with 1000 iterations for each trait separately. Significance thresholds were set above the 95% confidence intervals of the LOD score distribution. Third, kinship matrices and parental effects were calculated and multiparent QTL mapping was performed using the *selQTLMPP* function of *statgenMPP*. Parental effects are estimated for all families using a Hidden-Markov model framework called reconstructing ancestry blocks bit by bit (RABBIT). The effects are based on the hidden ancestral allele at each locus (Jung *et al*., 2025; Zheng *et al*., 2015).

### Determining physical position of QTLs on Norin 61

The physical position of each marker was known for Norin 61, since the SNP calling was performed using the Norin 61 reference genome. Hence, all significant markers within QTL were extracted along with their positions on the linkage map and corresponding physical position. Since *Rht-D1* is not annotated in Norin 61 and found on chromosome Un, its physical position was inferred using the genes that flank both sides of it as described in the method section. The physical QTL size was defined by the significant markers with the lowest and highest physical positions within a QTL. Two QTL were identical if their physical position was overlapping, or if the highest marker in one QTL was adjacent to the lowest marker of another QTL on the consensus map. Moreover, we classified detected QTL into a single QTL whenever overlapping QTL formed a continuous chain, even if some pairs did not overlap directly.

### Identifying candidate genes on Norin 61 reference genomes

To identify known regulators of pollen number or anther length on the reference genome of Norin 61, we performed a BLAST search and compared physical positions of already published QTL regions. We used BLAST+ (version: 2.16.0+) (Camacho *et al*., 2009) and performed a *tblastn* search of *A. thaliana RDP1* and *Zea mays CCT10* and *RPN1* on the Norin 61 reference genome. We used the genomic positions of *Ppd-D1* and *Rht-B1* on Norin 61 by Shimizu et al. (2021), as they were previously not annotated. Since *Rht-D1* was not annotated on the Norin 61 genome, we took genes up- and downstream (TraesCS4D03G0066900, TraesCS4D03G0066700, TraesCS4D03G0066500, TraesCS4D03G0068300, TraesCS4D03G0068700) of Chinese Spring *Rht-D1* (TraesCS4D03G0067100) and ran *blastn* on Norin 61. To determine the position of *Rht8,* marker *gwm261* was used to run *blastn* on Norin 61, as *Rht8* is linked to *gwm261* (Korzun *et al*., 1998). Next, we compared previously published QTL regions for anther length and pollen mass by performing *blastn* with the associated markers listed in the GWAS studies and blasted them against the reference genome of Norin 61.

### DNA extraction and genotyping *Rht-B1, Rht-D1,* and *Ppd-D1* in wheat diversity panel

We extracted DNA from the lines grown in the foil greenhouse to genotype the NAM parental lines and the small wheat diversity panel for *Rht-B1, Rht-D1* and *Ppd-D1.* Leaf tissue from adult plants was sampled, frozen in liquid nitrogen and then extracted by using the DNeasy Plant Mini kit (Qiagen, Cat. 69104) according to the manual. We used markers designed to amplify *Rht-B1a* (WT)*, Rht-B1b* (dwarf) and *Rht-D1b* (dwarf) (Ellis *et al*., 2002) and *Rht-D1a* (WT) (Guedira *et al*., 2010) (Suppl. Table 3). *Ppd-D1a* and *Ppd-D1b* alleles were genotyped based on the protocol of Beales et al. (2007) (Suppl. Table 3). Norin 10-NBRP (LPGKU2306), obtained through NBRP, carried only the mutant allele *Rht-B1b* (*Rht1*), while *Rht-D1* (*Rht2*) was found to be wild-type allele *Rht-D1a*. Although Norin 10 is known for the introduction of *Rht-B1b* and *Rht-D1b* during the green revolution, the difference may be attributed to the genetic variation within cultivars having the same name stored in different stock center, which are also documented in other species such as *Hordeum vulgare* (barley) or *A. thaliana* (Doyle *et al*., 2005; Dreiseitl and Zavřelová, 2022). From the landrace Dika neither *Ppd-D1a* or *Ppd-D1b* alleles were amplified despite repeated genotyping attempts.

We build linear mixed effect models to test if the allelic status of *Rht-B1b, Rht-D1b* and *Ppd-D1* can partially explain the difference in pollen numbers and anther length between landraces and cultivars. We used *Rht-B1, Rht-D1* and *Ppd-D1* allele status information as fixed effect in the model while genotype was treated as a random effect. Alternatively, we tested models including status (i.e., landrace, traditional, and modern cultivar), height, or days to flowering as fixed effects. Plant height was only available for season 2023. The best model was chosen after comparing the models with the Akaike Information Criterion (AIC) (Akaike, 1998). Models with differences in AIC lower than 2 were considered to be of equal fit. We calculated marginal and conditional *R^2^* for the models with the lowest AIC values (Nakagawa and Schielzeth, 2013).

## Results

### Pollen number and related traits of wheat landraces and cultivars

We grew 54 lines for two seasons in a foil greenhouse consisting of 23 landraces and 31 cultivars (12 traditional and 19 modern cultivars) originating from Europe, Asia, the Americas, Australia, and Africa to examine the variation in male reproductive traits within a wheat panel representing a subset of the global wheat diversity (Suppl. Table 1). In both seasons, anther lengths and pollen numbers were significantly higher in landraces than in cultivars (Figure 1A-F, H-I). Pollen diameter differed between landraces and cultivars significantly only in season 2022 (Figure 1G, J). Across both seasons, we observed extensive variation in anther lengths, pollen numbers, and pollen diameters (Suppl. Figure 2A-C for the season 2022, Suppl. Figure 3A-C for the season 2023). In 2023, pollen number ranged from 2895 (Nanking No. 25) to 10350 pollen grains per floret (CN2) (Suppl. Figure 3B, Suppl. Table 4, season 2023). Similarly, the anther lengths ranged from 3.22 (Claire) to 5.98 mm (CN2) (Suppl. Figure 3A, Suppl. Table 4, season 2023). Summarized data by each genotype for both seasons are shown in Suppl. Table 4.

**Figure 1:**
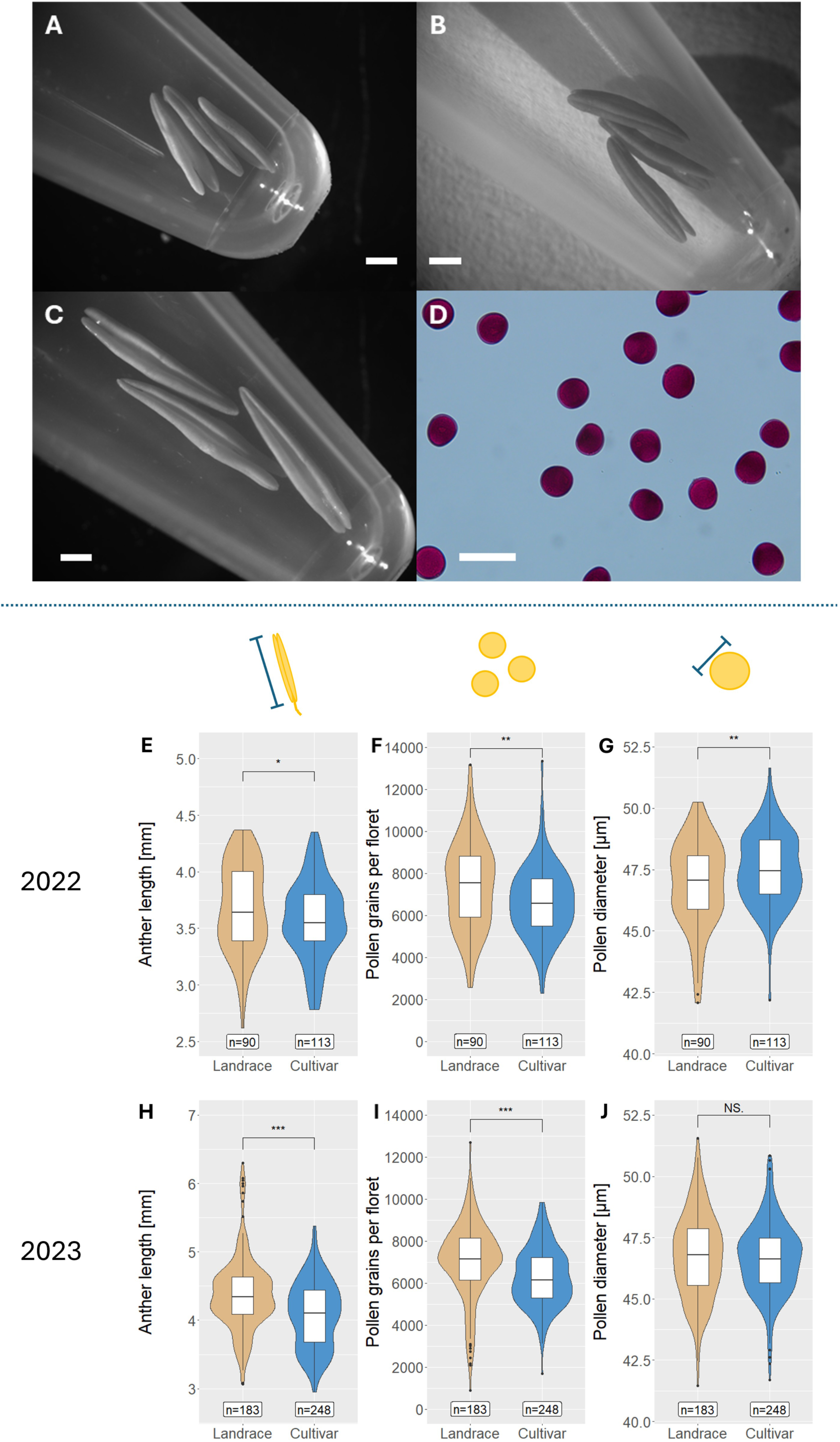
Variation of anther- and pollen-related traits among landraces and cultivars. Anthers of cultivars **(A)** Claire and **(B)** Norin 61 and landrace **(C)** CN2. **(D)** Norin 61 pollen grains stained by Alexander solution. White bars indicate **(A-C)** 1 mm and **(D)** 100 µm. Box- and violin plots showing data for **(D, G)** anther length, **(E, H)** pollen number, and **(F, I)** pollen diameter. **(D-F)** shows data for season 2022. **(G-I)** shows data for season 2023. Numbers written at the bottom of the panel indicate the number of measured florets (3 anthers) per category. Dots represent outliers. *** = p-value ≤ 0.001. ** = p-value ≤ 0.01. * = p-value ≤ 0.05. n.s. = not significant.

Based on the reduction of pollen number and anther length in cultivars, we examined the time course of the reduction by classifying cultivars into two categories: traditional (12 lines) and modern cultivars (19 lines; Suppl. Table 1). According to the definition of Balfourier et al. (2019), traditional cultivars are lines that were released before 1960 and the green revolution, while modern cultivars were released after 1960.

We found that modern cultivars produced a significantly smaller pollen number compared with landraces both in 2022 and in 2023 (Figure 2A, Suppl. Figure 4A). The same significant difference was also confirmed when the data in 2022 and 2023 were combined (Suppl. Figure 5A). Traditional cultivars had a significantly higher number of pollen grains than modern cultivars in 2023 and the combined dataset, although it was not significant in 2022, consistent with a lower sample number (Figure 2A, Suppl. Figures 4A, 5A). For anther length, modern cultivars showed significantly shorter anthers than landraces in 2023 and in the combined dataset (Figure 2C, Suppl. Figures 4C, 5C). In contrast to pollen numbers and anther lengths, pollen diameter did not show a significant difference between landraces and traditional cultivars in 2023 and in the combined dataset, while landraces showed significantly smaller pollen diameter compared to the traditional cultivars in season 2022 (Figure 2E, Suppl. Figures 4E, 5E).

**Figure 2:**
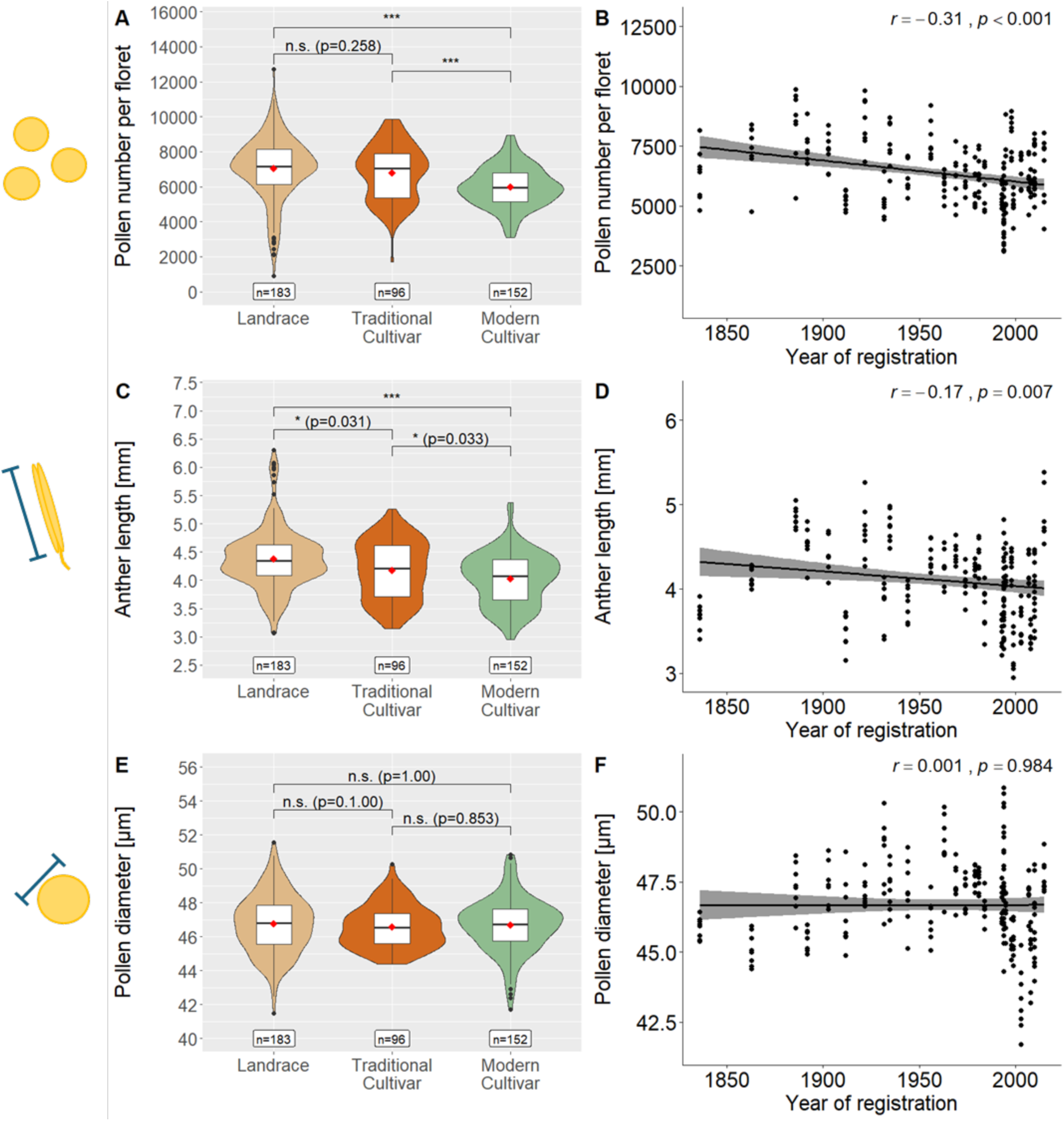
Comparison of **(A)** pollen number, **(C)** anther length and **(E)** pollen diameter for landraces, traditional cultivars and modern cultivars in season 2023. Pearson correlation between year of registration and **(B)** pollen number, **(D)** anther length, and **(F)** pollen diameter. n = number of analyzed florets (3 anthers). Red diamond indicates mean value. *** = p-value < 0.001. ** = p-value < 0.01. * = p-value ≤ 0.05. n.s. = not significant. Adjusted *P-values* calculated by Kruskal-Wallis-test and post-hoc Dunn’s test for **(A,C,E)**. Black dots represent outliers in **A,C,E**.

To further examine historical change of these traits, we plotted them with the years of registration, which is available for traditional and modern cultivars. In season 2023 we found negative correlations of pollen number (*r* = -0.31, *p-value* < 0.001) and anther length (*r* = -0.17, *p-value* = 0.007) with the year of registration, consistent with the reduction from traditional to modern cultivars (Figure 2B, D). In season 2022, the correlation was not significant, although negative trends were observed (pollen number *r* = -0.17, anther length *r* = -0.11) (Suppl. Figure 4B, D). In the combined dataset, significant negative correlations were detected (pollen number: *r* = -0.25, *p-value* < 0.001, and anther length: *r* = -0.15, *p-value* < 0.01) (Suppl. Figure 5B, D). Pollen diameter did not show a significant correlation with year of registration in any of the datasets (Figure 2F, Suppl. Figures 4F, 5F).

Broad-sense heritability estimates accounting for both seasons together were high for all three traits, with pollen number showing the highest value (*H^2^_Anther length_* = 0.66, *H^2^_Pollen number_* = 0.82, *H^2^_Pollen diameter_* = 0.66). Heritability estimates for each year separately were even higher, ranging from 0.89 for pollen number in season 2022 to 0.97 for anther length in season 2023 (Suppl. Table 5).

In both seasons, we calculated correlations among pollen-related traits (pollen number, anther length, and pollen diameter) and days to sampling, a proxy for flowering time (Suppl. Figure 6). Overall, correlations were often significant but low. Anther length and pollen number were positively correlated in both seasons, but the strength varied considerably (*r* = 0.32, *p-value* < 0.001 season 2022; *r* = 0.63, *p-value* < 0.001, season 2023). This suggests that while anther length is commonly used as proxy for pollen number, it may not always be a reliable indicator (see discussion). Pollen number and anther length were also positively correlated with days to sampling in both seasons (Suppl. Figure 6). Regarding pollen diameter, we observed positive correlations with anther length in both seasons (Suppl. Figure 6). In 2023, pollen diameter was negatively correlated with both pollen number and days to sampling; in contrast, no significant correlations were detected in 2022 (Suppl. Figure 6).

### Pollen number of NAM parental lines in chamber environments

Within the global collection of 54 wheat lines, 25 parental lines of the NBRP nested association mapping (NAM) population were grown to assess the variation of pollen-related traits among the NBRP-Wheat collection in Asia. These 25 lines comprised 14 landraces and 11 cultivars. We conducted two growth experiments under controlled growth chamber conditions.

In the first experiment, we confirmed the same pattern of higher pollen production in landraces under controlled condition, with a clearer difference between landraces and cultivars (Figure 3A, B, Suppl. Figures 7, 8). The median pollen number of each of the 14 landraces was larger than the median of any of the 10 cultivars (Figure 3B). Median pollen numbers per anther for each line showed a large variation, ranging from 917 (ZNK) to 2,820 (CN6). The common parent Norin 61 had a relatively low number, 1,300 pollen grains. The broad-sense heritability for pollen number in this experiment was high, further supporting its suitability for QTL mapping (*H^2^_Pollen number_* = 0.84) (Suppl. Table 5).

**Figure 3:**
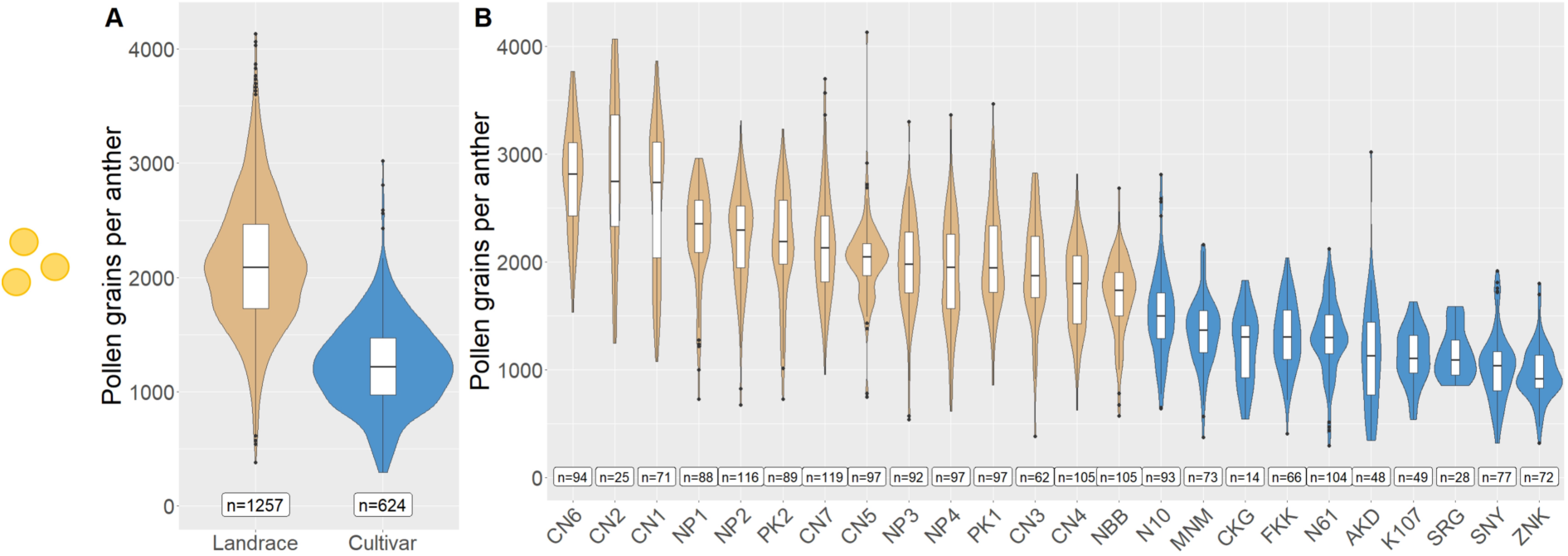
Comparison of pollen grains per anther between landraces and cultivars **(A)**, and among NAM parental lines **(B)** grown under controlled growth chamber environment. Parental line CGW is not included due to low sample number (see Suppl. Figure 7). The numbers written at the bottom of the panel indicate the number of analyzed florets (3 anthers). Dots represent outliers.

In the second growth experiment, four replicates per genotype stayed in the same environment (control) while four other replicates were moved to the heat treatment from the mid boot stage until the beginning of flowering. Consistent with the known negative effect of heat stress on wheat male germline development (Bheemanahalli *et al*., 2019; Shenoda *et al*., 2021; Xu *et al*., 2022), we found a 15% reduction of median pollen numbers averaged over all lines. This difference was driven by the reduction of pollen numbers in landraces, since cultivars in total showed no significant difference between control and heat treatment (Suppl. Figure 9A, B). Yet, when each cultivar is examined, a few cultivars such as K107 produced significantly less pollen grains with the heat treatment (Suppl. Figure 10). Although the landraces produced 18% less pollen grains under heat stress than under the control environments (Suppl. Figure 9B), they still showed an 11% higher median pollen number per anther than cultivars (Suppl. Figure 9C, D). Furthermore, heat stress significantly reduced pollen diameter in cultivars and landraces (Suppl. Figure 11A, B). This reduction was larger in landraces than cultivars (Suppl. Figures 10, 11, 12).

### Different and shared QTLs on pollen number and anther length detected in the NAM population

Based on the high heritability of pollen number, anther length and pollen diameter described above, we conducted QTL mapping using 1057 NAM lines composed of 13 RIL families in a field condition. These families were derived from the cross of 8 landraces and 5 cultivars with the common parental cultivar Norin 61 (Table 1).

Consistent with previous estimates, the heritability for all three measured traits was high in the NAM lines (*H^2^_Pollen number_* = 0.80, *H^2^_Anther length_* = 0.89, *H^2^_Pollen diameter_* = 0.84, Suppl. Table 5 and 6). The correlation between pollen number and anther length (0.68) was slightly higher than those in the foil greenhouses. There was more than a 4.5-fold difference in pollen numbers between the NAM RIL line with the lowest average pollen number per floret (KU-N04-022, 2100 pollen grains) and the two lines with the highest pollen number (KU-N06-036 and KU-N07-050, 9505 pollen grains) (Suppl. Table 6). Similarly, anther length ranged from 2.49mm (KU-N18-046) to 5.30mm (KU-N01-010) (Suppl. Table 6). While sampling date, as proxy for flowering date, showed a positive correlation with pollen number and anther length in the small wheat diversity panel, the two traits were negatively correlated in the NAM lines (Suppl. Figures 13-15). Based on these results, we tested different linear mixed effect models to calculate BLUP values for anther length and pollen number for later use in multiparent QTL mapping. All tested models included genotype as random factor. The best model for pollen number included days to sampling, while that for anther length included the days to sampling and pollen diameter as fixed factors.

Multiparent QTL mapping using the nested association mapping design was performed using pollen number, pollen diameter and anther length, as well as BLUP values calculated for pollen number and anther length. When we used the non-adjusted phenotypes as input, a significant QTL on chromosome 5B for pollen number (named *QPN.uzh-5B.1)* and four QTLs for anther length (similarly named *QAN.uzh*) on chromosomes 1A, 2D, 4D, and 5B were detected (Figure 4A, Suppl. Figure 16). More significant QTLs were detected when BLUP values were used as phenotype input: two additional QTL peaks for pollen number on chromosomes 1A and 2D (Figure 4B, Suppl. Figures 16, 17) and two additional QTLs on chromosomes 1B and 4B for anther length. Some of the QTL peaks, such as those on chromosome 5B, were significant both for pollen number and anther length (Table 2, Suppl. Figures 16F, 17F). Importantly, we detected a QTL significant only for pollen number *QPN.uzh-1A.1* on chromosome 1A (Table 2, Figure 4B, Suppl. Figures 16A, 17A). Across traits, individual significant QTLs explained between 3.62% and 15.77% of the phenotypic variation (Table 2, Suppl. Figure 18). QTLs detected based on BLUP values explained 34.55% of the phenotypic variation for anther length and 25.07% for pollen number. No significant QTL of pollen diameter was detected (Suppl. Figure 19)

**Figure 4:**
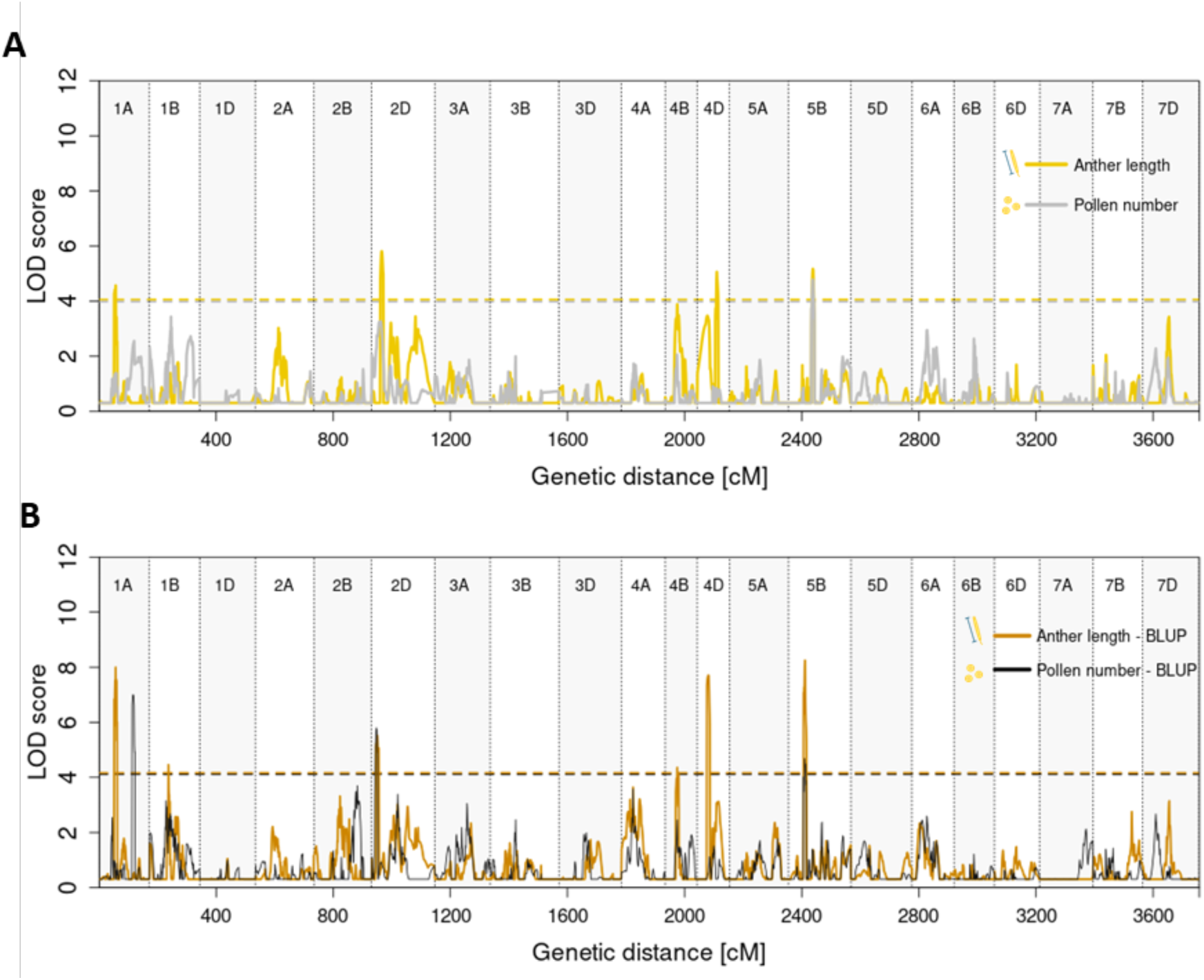
Multiparent QTL mapping for pollen number and anther length using **(A)** measured phenotype as input and **(B)** BLUP values. BLUP values were calculated using genotype, replicate and days until sampling as random effects. Model for anther length BLUP calculation additionally included pollen diameter. Dashed lines indicate significance threshold.

**Table 2:** Summary table for all detected QTL for anther length and pollen number. PVE = phenotypic variation explained.

| QTL name | Phenotype | Position (cM) | Physical Position | LOD score | PVE | QTL size on physical map (Mb) | comments |
| --- | --- | --- | --- | --- | --- | --- | --- |
| *QAL.uzh-1A.1* | Anther length | 53.178 - 61.510 | 59494715 - 446179078 | 4.57 | 5.07% | 386.68 | Considered the same QTL |
| *QAL.uzh-1A.1* | Anther length - BLUP | 53.718 - 61.510 | 59494715 - 446179078 | 8.00 | 7.79% |  |  |
| *QAL.uzh-1B.1* | Anther length - BLUP | 57.692 - 69.246 | 421132112 - 544903148 | 4.46 | 6.11% | 123.77 |  |
| *QAL.uzh-2D.1* | Anther length | 28.184 - 39.338 | 26183491 - 35514780 | 5.81 | 11.13 % | 23.13 | Considered the same QTL as the lowest and highest markers are adjacent. Overlap with *QPN.uzh-2D.1* |
| *QAL.uzh-2D.1* | Anther length - BLUP | 19.281 - 22.192 | 12200453 - 22878690 | 5.51 | 8.36% |  |  |
| *QAL.uzh-4B.1* | Anther length – BLUP | 36.722 - 46.520 | 24685058 - 354814806 | 4.36 | 3.62% | 330.13 |  |
| *QAL.uzh-4D.1* | Anther length | 62.581 - 70.659 | 397765787 - 465963176 | 5.06 | 9.13% | 68.20 |  |
| *QAL.uzh-4D.2* | Anther length - BLUP | 28.872 - 42.363 | 13618714 - 27337800 | 7.71 | 3.55% | 13.72 |  |
| ***QAL.uzh-5B.1*** | Anther length | 77.883 - 88.710 | 516679253 - 538592288 | 5.17 | 5.78% | 21.91 | Overlap with *QPN.uzh-5B.1* |
| ***QAL.uzh-5B.2*** | Anther length - BLUP | 50.945 - 60.767 | 381145912 - 465279960 | 8.25 | 5.12% | 84.13 | Overlap with *QPN.uzh-5B.2* |
| ***QPN.uzh-1A.1*** | Pollen number - BLUP | 116.608 - 120.969 | 554063961 - 559128674 | 6.99 | 5.19% | 5.06 |  |
| ***QPN.uzh-2D.1*** | Pollen number – BLUP | 8.611 - 20.566 | 12200453 - 28343554 | 5.80 | 15.77 % | 23.13 | Overlap with *QAL.uzh-2D.1* |
| ***QPN.uzh-5B.1*** | Pollen number | 77.883 - 88.710 | 516464717 - 538592288 | 4.35 | 3.90% | 21.91 | Overlap with *QAL.uzh-5B.1* |
| ***QPN.uzh-5B.2*** | Pollen number - BLUP | 49.917 - 59.725 | 370290486 - 465279960 | 4.68 | 4.11% | 84.13 | Overlap with *QAL.uzh-5B.2* |

Within the genomic interval of QTLs on chromosomes 4B (*QAL.uzh-4B.1*) and 4D (*QAL.uzh-4D.2*), we found the two green revolution genes *Rht-B1* and *Rht-D1,* respectively (Table 2, Suppl. Tables 7 and 8). On chromosome 2D, the genomic regions overlapping with *QPN.uzh-2D.1* and *QAL.uzh-2D.1* encompassed *Ppd-D1* and *Rht8.* Other QTLs detected in the study neither overlapped with known GWAS peaks for anther length or pollen mass (Suppl. Table 9; Boeven et al. 2016, Song et al. 2018), nor contained homologs of known regulators of pollen number in other species (i.e. *AtRDP1, ZmRPN1, ZmCCT10*, Suppl. Table 8).

We further examined the QTL effects in each parental line. Here, we mainly focus on the effects of Norin 61 as it showed significant values for all QTL, consistent with the highest sample number as the common male parent in the NAM population (Suppl. Figure 18). Norin 61 showed a positive effect for *QAL.uzh-4B.1,* while the effect was significantly negative for *QAL.uzh-4D.2*. These effects would be in line with Norin 61’s *Rht-B1a* and *Rht-D1b* alleles and their known influence on anther length (Suppl. Figure 18C, Suppl. Table 10; Schierenbeck et al. 2024). At the anther length locus, *QAL.uzh-1A.1*, two landraces (NP4, PK1) and a cultivar (N10) had an allele associated with a significant increase, in contrast to the negative effect observed in Norin 61 and Chinese Spring (Suppl. Figure 18C). Although the phenotypic variation explained by the loci is relatively small, the effect sizes suggest that alleles with both positive and negative impacts are segregating in Norin 61 and potentially in other parental lines.

### Small but significant contribution of the genomic region encompassing green revolution genes *Rht-B1, Rht-D1* and *Ppd-D1* to the reduction of pollen number

The categorization of modern cultivars is based on the green revolution and the accompanied introduction of mutant *Rht* alleles as well as *Ppd-D1a* that is linked to *Rht8* in the global breeding pool. To examine whether *Rht-B1* and *Rht-D1* have an influence on the reduction of pollen number and anther length in our samples, we genotyped all the lines using published markers (Guedira et al., 2010; Ellis et al., 2002, Suppl. Table 3). Among the 54 lines, 11 carried the *Rht-B1b* allele, 8 carried the *Rht-D1b* allele, while none had both. The remaining lines carried the wild-type alleles (*Rht-B1a* and *Rht-D1a*) (Suppl. Table 10). For *Ppd-D1,* 15 lines carried *Ppd-D1a,* including all Japanese cultivars. The remaining lines had the wild-type allele *Ppd-D1b* (Suppl. Table 10). Lines carrying mutant alleles *Rht-B1b, Rht-D1b* or *Ppd-D1a* had significantly lower pollen numbers and shorter anthers compared with lines carrying wild-type alleles (Suppl. Figures 20-22). This supports our QTL mapping results in which peaks overlapped with the *Rht* genes and *Ppd-D1* (Suppl. Figure 17C-E).

We fitted a series of linear mixed-effects models to assess the effects of *group* (landrace, traditional or modern cultivar), *Rht-B1 and Rht-D1* allele status, *Ppd-D1* allele status, plant height, and flowering time on pollen production per floret (Suppl. Table 11). All models included line as a random effect to account for variation in pollen number among genotypes. Model comparisons based on Akaike Information Criterion (AIC) revealed that the full model including all fixed effects (group, *Rht-B1* and *Rht-D1* allele status, *Ppd-D1* allele status, days to flowering, and height (only for season 2023)) had the lowest AIC, indicating the best fit (model_8 (season 2022) and model_16 (season 2023) in Suppl. Table 11). Based on calculation of marginal *R^2^*, fixed effects explained 14.41% of the variance in season 2023 while line as random effect together with fixed effects explained 71.93% of the variation (conditional *R^2^*). In season 2022 marginal *R^2^* was 10.20% and conditional *R^2^* was 69.11% (Suppl. Table 11). Simpler models with only a single fixed effect (e.g. dwarfing allele status or group) showed higher AIC values (>2), suggesting that multiple traits contribute to variation in pollen number. The relatively small contribution of green revolution genes to pollen number is consistent with the multiparent QTL mapping described above. Among the three loci, only the QTL region encompassing *Ppd-D1*-*Rht8* was significant for the BLUP values of pollen number and explained 15.77% of the variation. Considering the BLUP values of anther length, the QTL regions encompassing *Rht-B1, Rht-D1*, *Ppd-D1-Rht8* explained 3.62% (*QAL.uzh-4B.1*), 3.55% (*QAL.uzh-4D.2*), and 8.36% (*QAL.uzh-2D.1*) of the variation, respectively.

## Discussion

### Reduction of pollen number in modern cultivars

The reduced number of pollen grains is a hallmark of the selfing syndrome (Shimizu and Tsuchimatsu, 2015; Sicard and Lenhard, 2011; Tsuchimatsu and Fujii, 2022). Here, we found that the bread wheat cultivars used in this study have a reduced pollen number and anther length compared with landraces. By analyzing cultivars registered between 1836 and 2015, we found that modern cultivars (post-1960) produce fewer pollen grains than traditional cultivars (pre-1960). This was also supported by the negative correlation between pollen number and the year of registration. A recent and rapid reduction of anther extrusion, which is another characteristic trait of selfing syndrome, was also reported in bread wheat (Boeven *et al*., 2016). Together, these studies show that two traits of the selfing syndrome reflect evolution in action in bread wheat during the past 200 years of breeding.

In crop species, increased selfing rate and selfing syndrome often evolved during domestication and subsequent breeding history from landraces to cultivars (Akagi *et al*., 2022; Dempewolf *et al*., 2012). Domesticated rice showed higher selfing rate and shorter anthers than wild progenitor species, indicating reduction of pollen number during domestication (Oka and Morishima, 1967). Landraces are often characterized by genetic heterogeneity suggesting outcrossing exemplified by studies of sorghum and tetraploid wheat (Barnaud *et al*., 2008; Tsegaye, 1996). The landraces of bread wheat used in this study may show a higher outcrossing rate, which should be verified in their original habitats.

The shift from landraces and traditional cultivars to modern cultivars during the green revolution is characterized by the introduction of semi-dwarfing *Rht* alleles and photoperiod-insensitive *Ppd* alleles (Balfourier *et al*., 2019; Beales *et al*., 2007). Our data supported their contribution to the variation in pollen number and anther length as shown by previous studies (Okada *et al*., 2019; Schierenbeck *et al*., 2024), but their contributions are relatively small in the NAM lines (3.62%, 3.55% and 8.36% of the phenotypic variation in BLUP of anther length by the genomic region encompassing *Rht-B1, Rht-D1* and *Ppd-D1* or *Rht8*, respectively; 15.77% of the phenotypic variation of BLUP of pollen number by the region encompassing *Ppd-D1* or *Rht8*). The data suggest that, in addition to green revolution genes, many QTLs contributed to the reduction in pollen number over the past 200 years.

The selective force driving the evolution of selfing syndrome, including reduced pollen number, has long been discussed (Shimizu and Tsuchimatsu, 2015; Tsuchimatsu and Fujii, 2022). On the one hand, variants decreasing viable male gamete numbers are deleterious by decreasing reproductive success unless there are pleiotropic effects (Willis, 1999). The spread and fixation of such deleterious variants can be enhanced when the effective population size is small due to an increased selfing rate or due to bottlenecks during domestication and breeding history in crop species. In bread wheat, natural genetic diversity diminished from landraces to modern cultivars (Balfourier *et al*., 2019; Tanksley and McCouch, 1997). On the other hand, evolutionary studies suggested that reduced investment in male fertility can increase seed mass and female fertility due to a postulated tradeoff and thus can be advantageous (Charnov, 1982; Tsuchimatsu and Fujii, 2022). In wild species *A. thaliana*, the allele of *RDP1* conferring reduced pollen number showed signature of selection, suggesting that the reduced pollen grain was adaptive (Tsuchimatsu *et al*., 2020). In bread wheat, a study using 10 cultivars detected no significant correlation between the number of pollen grains and yield (Chowdhry *et al*., 1992). A larger scale measurement would be valuable to test if the reduction of pollen number can confer increased yield in bread wheat.

Recently, selection for larger pollen and larger anthers during domestication have been described in rye (Waesch *et al*., 2025). In our study we did not detect any significant differences between landraces and cultivars in pollen diameter (Figure 2C). This difference is consistent with contrasting outcrossing rates: predominantly outcrossing rye and predominantly selfing wheat (Burgarella *et al*., 2024). The increased pollen size in rye might be an adaptation to change in its ecology, as cultivated rye is planted in higher densities compared to feral rye. Feral rye needs to transport its pollen farther distances, while smaller pollen facilitates the distribution distance (Waesch *et al*., 2025).

### Different QTLs underlying pollen number and anther length

Anther length is correlated with pollen number and has therefore been widely used as a proxy of male fertility in functional, evolutionary and agricultural studies because counting pollen number was traditionally too tedious for large-scale phenotyping (Kakui *et al*., 2020a; Langer *et al*., 2014). Pollen number has a direct link to male fertility and fitness in contrast to anther length, although the latter may affect flower morphology such as anther extrusion (Boeven *et al*., 2016). We recently established a pollen-counting method that is efficient enough for mapping populations (Kakui *et al*., 2020a; Kakui *et al*., 2020b). We first found that the correlation between pollen number and anther length are not very high. Our data ranged from low to moderate correlations between the two traits in two foil greenhouse seasons (*r* = 0.32 and 0.63, Suppl. Figure 6 A,B) and NAM lines (*r =* 0.68). Previous studies reported also a wide range of correlation from moderate (*r* = 0.41) (Chowdhry *et al*., 1992) to high correlation coefficients in bread wheat (*r* = 0.87-0.93) (De Vries, 1974; Milohnic *et al*., 1976; Nguyen *et al*., 2015). These data suggests that the correlation may not be very high, depending on cultivars and growth conditions. It is also plausible that pollen number is affected by developmental constraints such as the three-dimensional structure of anthers and the resulting balance of cell division and elongation.

Next, using more than 1000 NAM lines, we found that QTLs detected for pollen number overlapped only partially with those for anther length. Most importantly, *QPN.uzh-1A.1* on chromosome 1A was detected only by QTL mapping of pollen number and not anther length. The mapped interval (5.06 Mb) for *QPN.uzh-1A.1* was the smallest among all QTLs detected in this study, and it is a promising QTL of future fine mapping and marker-assisted breeding. In our QTL mapping, we also found QTLs that are significant for anther length but not for pollen number. However, for two regions, which encompass *Rht-B1b* on chromosome 4B and *Rht-D1b* on chromosome 4D, the QTLs also have small but significant effects on pollen number that were detectable by additional genotyping. It is possible *Rht-B1* and *Rht-D1* genes, which encode DELLA proteins, primarily have an observable effect on cell elongation of anthers rather than on cell division of pollen lineages (Cheng *et al*., 2004; Schierenbeck *et al*., 2024).

### Pollen number affected by phenology and heat stress

The relationship between fertility and phenology has been studied in natural and crop species in the currently changing world (Hedhly *et al*., 2009; Liu *et al*., 2020; Shimizu *et al*., 2011). Here, we found a significant relationship between days to flowering (using days to sampling as a proxy) and pollen number as well as with anther length. However, the direction of the correlation differed between experiments, with positive correlations using the 54 cultivars grown in the foil greenhouse and negative correlations using the 1057 NAM lines in the field. These results are in line with the study of Komaki and Tsunewaki (1981), showing a quadratic relationship between anther length and days to flowering. Anther length in that study was longest in cultivars that flowered at the intermediate time points during the season, while both early- and late-flowering cultivars produced shorter anthers. Our study also suggests that there is an optimum time to produce a high number of pollen grains, and deviations from this optimum can reduce pollen number due to developmental and environmental conditions. As a developmental constraint, early flowering cultivars may not have accumulated adequate resources to produce many pollen grains. Low and high temperatures in the early and late seasons, respectively, might reduce pollen number. Here, we showed that a relatively moderate heat treatment reduced the pollen numbers of landraces used in this study. It is possible that the NAM population showed a negative correlation consistent with its growth condition in the field in Kyoto, in which the temperature toward the end of the growth season could have negatively affected the pollen number.

The relevance of phenology on pollen number and anther length is highlighted by maize *CCT10*, which was originally identified as a flowering regulator and was shown to affect pollen number (Li et al., 2022; Liu et al., 2020). The connection is further supported in bread wheat by the contribution to anther length and pollen number of the genomic regions encompassing *Ppd-D1,* encoding a pseudoresponse regulator protein regulating heading and flowering time (Okada *et al*., 2019) (Suppl. Figures 17C, 21). The association of heading-time genes and pollen number may be explained by three hypotheses that are not mutually exclusive. First, it is conceivable that *Ppd-D1* or maize *CCT10* affected flowering time and thus the environment at anther development, leading to the change in pollen number. Second, heading-time genes might have a pleiotropic effect directly on stamen development separately from flowering time. Third, neighboring genes, such as *Rht8* linked to *Ppd-D1,* can affect pollen number.

Heat stress is a major threat in many wheat growing regions worldwide and increasingly so in the coming years due to climate change (Asseng *et al*., 2015; Zhao *et al*., 2017). Many studies showed low pollen viability by heat stress (Chaturvedi *et al*., 2021; Harsant *et al*., 2013). Here we found a reduction in pollen number for lines that experienced heat stress from mid-boot stage until flowering (Suppl. Figure 9). This is consistent with previous studies reporting reduced anther length in bread wheat (Xu *et al*., 2022) and reduction in pollen number in peanut, tomato and *Brassica napus* after heat stress (Prasad *et al*., 1999; Pressman *et al*., 2002; Rosenberger *et al*., 2024). This reduction was mainly driven by landraces, as cultivars in total did not show a significant difference. The stable pollen number of Japanese cultivars in the NAM parental lines may be attributed to unconscious selection for heat tolerance in the hot Japanese environments or to the loss of high phenotypic plasticity observed in landraces. Yet, when each cultivar is analyzed separately, a few cultivars suffered from the heat treatment producing reduced numbers of pollen grains, which may be mitigated by breeding utilizing landraces.

### Pollen number in hybrid breeding and in changing climates

Our identification of QTLs underlying pollen number is a critical step to increase or decrease pollen number in breeding. As discussed above, decreasing pollen number can potentially increase yield components postulated by sex allocation theory (Charnov, 1982). In turn, increasing pollen number can be advantageous in facilitating hybrid breeding. Here, we showed that bread wheat exhibits a large variation in pollen number, with more than a three-fold difference among the foil greenhouse lines in 2022 and 2023 (Suppl. Figures 2B, 3B, Suppl. Table 4) and more than a four-fold difference between NAM lines with lowest and highest pollen number (Suppl. Table 6). Together with high heritability, our findings will facilitate the establishment of genotypes with high pollen number. Although modern cultivars tend to show reduced pollen number, our results highlight that traditional cultivars and landraces are promising resources to increase pollen number. Considering a high variation in pollen number and high heritability in the wild selfing species *A. thaliana* with 4-fold variation among natural accessions (Tsuchimatsu *et al*., 2020), we suggest that high heritable variation of pollen number in selfing species can be segregating both in natural and crop selfing species. Previous studies showed the effect of green revolution gene *Rht-B1* and *Rht-D1* on anther length (Okada *et al*., 2019), while the genotyping data here also indicates an negative effect on pollen number (Suppl. Figure 20). Still, a constraint in using these loci to modify pollen number is their pleiotropic effect on organ size, such as stem height that potentially affects pollination efficiency (Schierenbeck *et al*., 2024). Here, we identified new loci affecting pollen number such as *QPN.uzh-1A.1* which are located outside of genomic regions encompassing green revolution genes, and propose that they will be useful to control pollen number in the future.

Interest in assessing pollen fertility in breeding has increased in recent years (Heidmann and Di Berardino, 2017; Impe *et al*., 2020; Langedijk *et al*., 2023). For example, many hazelnut cultivars showed lower pollen viability compared with wild plants (Ascari *et al*., 2020), suggesting that male fertility has been reduced during breeding. Our study demonstrated that pollen numbers are highly heritable, and that some individuals in the mapping population produce low pollen numbers. Routine evaluation of pollen fertility using available phenotyping methods could therefore help to avoid the selection of cultivars with reduced male fertility in breeding programs. We should emphasize that most of the variance of pollen number is not explained by the identified QTLs despite high heritability, suggesting polygenic architecture. This underlines the potential of identifying more QTL and the potential application of genomic selection using increased sample numbers.

## Author contributions

NBH, HK, SN and KKS designed the study. MN created the NAM population. NBH, HK, MO performed the experiments and collected the data. NBH, HK, MO, KJ, JN analyzed the data with the help of TW, BK, SN, KKS. Seeds for the NAM parental lines and the NAM RIL families were provided by MN and SN. NBH wrote the paper with inputs from HK, MO, KJ, KKS, TW, SN, BK. All authors discussed and commented on the manuscript.

## Acknowledgments

We thank Dr. Reiko Akiyma and Dr. Yasuhiro Sato for statistical advice, Marcel Brasser for technical help in setting up the foil house experiments and Aki Morishima for support in wet lab experiments. This study was funded by URPP Evolution in Action of the University of Zurich to KKS, TW and BK, the European Union’s Horizon2020 research and innovation program under the Marie Sklodowska-Curie (MSC) grant agreement No 847585 to KKS and SN, UZH Global Strategy and Partnerships Funding Scheme of the University of Zurich to KKS, SN and BK, JSPS International Leading Research Grant Number [JP22K21352] and JST CREST JPMJCR16031 to KKS and SN, Swiss National Science Foundation [310030_212551 and 31003A_1823181], JSPS KAKENHI Grant Numbers [JP21H05366, JP22H02316, JP23K23582 and JP22H05179] to KKS, JSPS KAKENHI Grant Numbers [JP22H02307, JP22K19172] to HK, and MEXT-NBRP-wheat, Genome Information Updating Program to SN and KKS.

Seeds of NAM parental lines and NAM families were provided by the National BioResource Project-Wheat with support in part by the National BioResource Project of the MEXT, Japan.

We thank the John Innes Centre Germplasm Resources Unit, a National Bioscience Research Infrastructure supported by the UKRI-BBSRC, grant number BBS/E/JI/23NB0001 for conserving and supplying germplasm through www.seedstor.ac.uk and the INRA Small Grain Cereals Biological Resources Centre for providing seeds of wheat landraces and cultivars.

